# Population-level encoding of somatosensation in mouse sensorimotor cortex

**DOI:** 10.1101/2025.01.21.634118

**Authors:** Seungbin Park, Megan H. Lipton, Maria C. Dadarlat

## Abstract

Somatosensation constructs the body’s dynamic sense of state and allows for dexterous and precise movements. The heterogeneous responses of single neurons in sensorimotor cortex to somatosensation have led to disparate views of the computational role of this brain area in sensori-motor processing. Here, we use population-level analyses of neural activity recorded during passive limb movements to assess the structure and to summarize the properties of neural encoding in sensorimotor cortex. We used 2-photon imaging to record the activity of thousands of neurons in eight anesthetized mice during passive deflections of each limb. We additionally analyzed neural responses to passive limb movements in eight awake mice, sourced from an open dataset [1]. We employed principal component analysis on the neural activity in each dataset and found that a small fraction of principal components explained a large fraction of variance in the neural responses. Low-dimensional representations of limb movements were well conserved across animals, including the orthogonal representations of ipsilateral and contralateral limbs. This organization of somatosensory information mirrors the well-known structure of neural encoding of motor commands in sensorimotor cortex. Furthermore, neural populations dually encoded both changes in joint angles during movements and more abstract information, i.e., the direction of limb movement. Increasing the size of the neural population improved encoding of both types of movement information and better differentiated representations from movements in opposing directions. Together, these results demonstrate that population-level encoding of somatosensory information in mouse sensorimotor cortex is structured to facilitate sensorimotor integration across the brain.

## INTRODUCTION

Somatosensation, which includes both proprioception and touch, is essential for the control of precise movements [2–5]. Proprioception, the sense of the body’s position and movement through space, is required to coordinate complex multi-joint and multi-limb movements, such as buttoning a shirt, tying a shoelace, and playing an instrument – where visual feedback alone is insufficient for precision and accuracy [6–11]. Tactile information similarly guides fine object manipulation, providing information about contact location, pressure, and timing [3, 4, 12].

The neural basis of somatosensation may be studied by recording neural activity during externally-imposed perturbations (passive limb movements) in which the subject does not actively control its limb position [1, 5, 13, 14]. Within the primary somatosensory cortex, neurons that respond to both the direction and amplitude of limb movements have been identified [1, 15, 16]. However, neural activity also correlates strongly with other movement variables, such as limb position, speed, movement amplitude, and the phase of the limb stance/swing cycle during locomotion [1, 15, 17–19]. Such variability in neural responses across behavioral tasks supports the argument that neurons are in fact encoding whole-limb kinematics, including the configurations of multiple joints in the arm and hand – information that is derived from the activity of muscle spindles in the limb [13, 20–22]. However, even within a fixed behavioral task, there is significant variability in the responses of single neurons within a population, prohibiting a conclusive statement regarding the “information” encoded by somatosensory cortex. Addressing this question is important to better understand the neural mechanisms involved in sensory-guided movements.

Instead of examining single neurons, we propose to “summarize” the information encoded by the sensorimotor cortex by using population-level analysis of neural activity patterns [23, 24]. Population-level neural activity can be captured by projecting the activity of neurons into a lower-dimensional space via some form of dimensionality reduction [25, 26]. Analyzing neural activity patterns during behavior in this low-dimensional “state space” has been pivotal for understanding cortical encoding of vision, olfaction, motor planning, and active sensorimotor processing [25, 27–32]. In the mouse, somatosensory feedback from the forelimb is distributed across both somatosensory and motor cortices [1]. We therefore hypothesized that analyzing the low-dimensional representations of somatosensation in mouse sensorimotor cortex may elucidate the neural structure and function of these brain areas in processing somatosensation during passive limb movements.

Analyzing the low-dimensional representation of neural encoding of somatosensation also facilitates a direct comparison to the known structure of neural encoding of motor commands [30]. From the traditional perspective of sensorimotor integration, sensory and motor signals must be combined for sensory information to guide movements. One way to achieve such integration is by converting sensory and motor signals into a common reference frame [33]. Applying a state-space perspective to neural encoding of somatosensation offers a rather literal way to compare these reference frames [34]. For example, neural population activity within motor cortex during reaching is well described as a low-dimensional dynamical system, where the rate of change of a neural population’s activity is a function of ongoing neural activity and external input [17, 30, 35, 36]. However, the relative dimensionality of somatosensory information in these brain areas is unknown. Another example is that state-space encoding of ipsilateral and contralateral limbs during active movements are orthogonal in motor cortex [37–39]. While sensorimotor cortex shows clear responses to ipsilateral limbs, the activity patterns of single neurons responding to ipsilateral limb movements are highly variable [40]. Analyzing the low-dimensional representation of ipsilateral and contralateral limbs in mouse sensorimotor cortex may clarify their computational role in this brain area. Directly addressing these questions will yield insight into the neural processes underlying sensorimotor integration – the closed-loop process by which humans and animals use sensory information to guide movements.

In this manuscript we use principal component analysis (PCA) to “summarize” activity within a large population of neurons recorded from mouse sensorimotor cortex in both anesthetized and awake mice. Using low-dimensional representations, we found that the structure of neural activity in mouse sensorimotor cortex that arises from somatosensory inputs from the limbs mirrors the observed low-dimensional structure of motor commands. These representations were well-conserved across animals. Furthermore, we demonstrated that correlated activity across neural populations encodes sensory variables that generalize single neuron representations. Together, our results illustrate how the parallel structure of somatosensation and motor commands may facilitate sensorimotor integration across the brain.

## RESULTS

In this study, we examined population-level neural encoding of limb movements in mouse sensorimotor cortex in two contexts: 1) during passive deflections of each of the four limbs of anesthetized mice (Figure 1A), and 2) during passive movements of a single forelimb contralateral to the cortical hemisphere in which neural signals were recorded (using an open-source dataset [1]; Figure 1B). These complementary datasets each offer unique insights into neural encoding of somatosensation.

**Figure 1:**
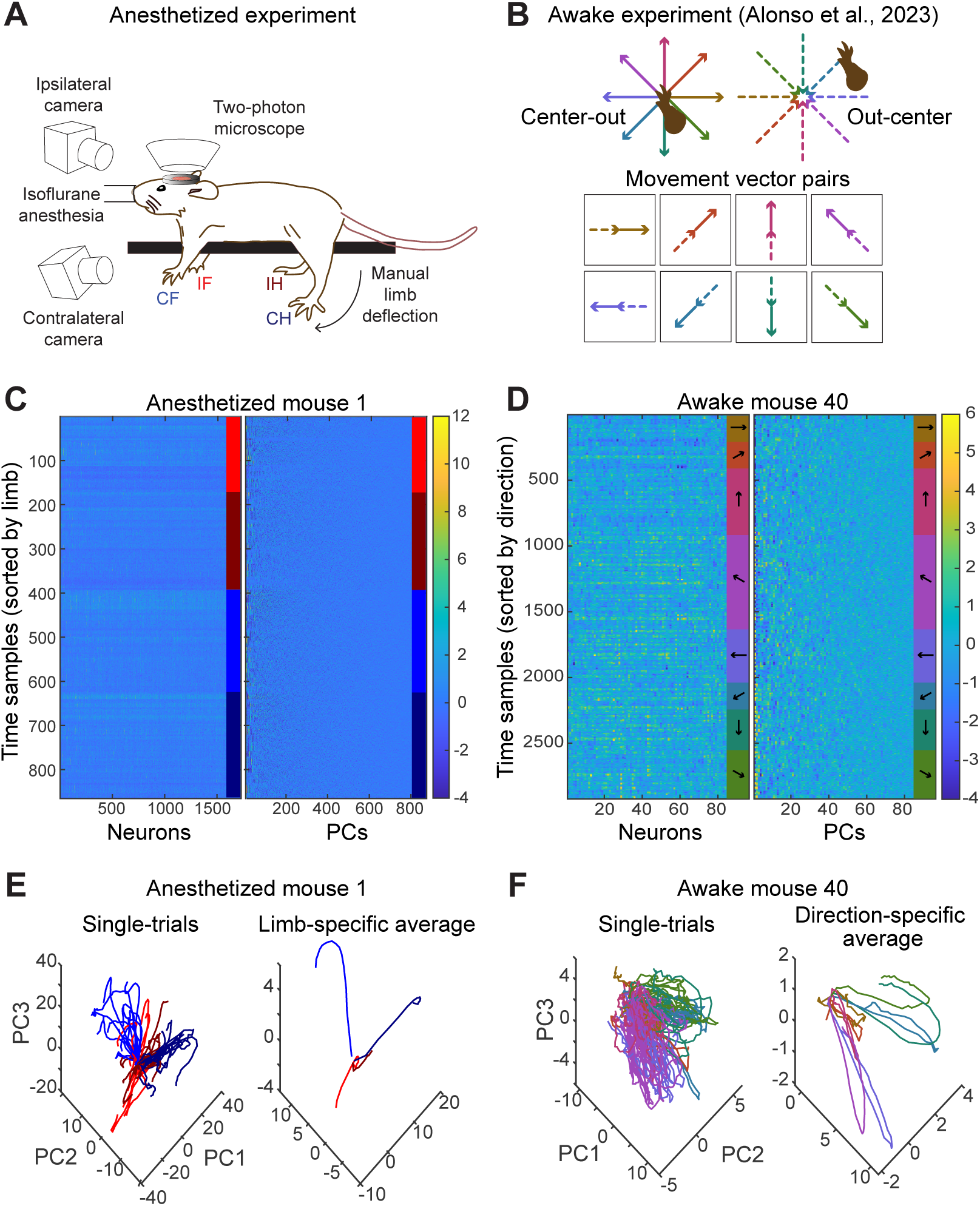
Experimental and analytical overview. **A** Experimental setup for recording neural responses to limb deflection in anesthetized mice. **B** Schematic showing center-out and out-center passive planar movements in [1]. Notably, the same movement vectors are repeated with different start and end points in the two conditions. **C** A comparison of single-neuron and PCA representations of population-level activity during limb deflection in a single anesthetized mouse. Z-scored responses for single neurons (left) and in PCA space (right) to limb deflection, shown for each time sample of each trial, with trials sorted by limb. Limb colors as in (A). **D** As in (C), for a single awake mouse. Colors indicate movement directions as in (B). **E** Low-dimensional representation of time-varying neural activity in an anesthetized mouse during passive limb deflections, shown both for individual trials and trial averages for each limb. Data comes from the same animal as in (C). **F** Low-dimensional representation of time-varying neural activity in an awake mouse during passive limb deflection, shown both for individual trials and trial averages for each limb. Data comes from the same animal as in (D).

### Low-dimensional encoding of limb movements in mouse sensorimotor cortex

How are neurons in the sensorimotor cortex encoding limb movements as a population? One way to address this question is to calculate the dimensionality of neural activity within mouse sensorimotor cortex. The dimensionality of neural responses identifies the size of the subspace defined by observed neural activity patterns, which may be much lower than the total number of neurons in the population [34]. Given the low dimensionality of activity patterns in somatosensory thalamus and motor cortex during active movements [25, 30], a straightforward prediction is that encoding of somatosensation in sensorimotor cortex is similarly low-dimensional. That is, neural activity would be dominated by relatively few components (e.g., [41]). Alternatively, the significant anatomical expansion from the thalamus to cortex could mediate a corresponding increase in the dimensionality of somatosensory encoding that serves to enhance the discriminability of stimuli [42–44].

To test this idea, we found the principal components of the neural responses to passive limb movements. Plotting the neural response matrix in PC space clearly shows that neural activity is highly correlated and concentrated in a fraction of the total possible dimensions (Figure 1C,D). We observed distinct neural activity patterns associated with passive movements of each limb and with movements of a single limb to different directions (Figure 1E,F). To calculate the dimensionality of neural activity, we found the number of significant PCs within the datasets. A PC was defined as being significant if its associated eigenvalue was larger than the threshold set by the Marchenko-Pastur law, which calculates the largest eigenvalue that can be found by a random matrix of a particular size [45, 46]. To compare across datasets composed of different numbers of neurons, as well as with past work, the number of significant PCs was then normalized by the total number of independent variables [47, 48]. Based on this analysis, we find that while the number of significant PCs made up a relatively small fraction of the total number of variables (neurons, below 0.15), they explained a substantial fraction of the total variance in the dataset (0.2–0.7 cumulative proportion of variance explained; Figure 2A,B and Tables S1,2). The ratio of the fraction of significant components to cumulative variance explained is highly conserved both across animals and across the anesthetized and awake datasets, despite both the difference in task structure in the datasets and the variability in the number of neurons recorded in each animal. Furthermore, these values are consistent with past work evaluating the dimensionality of neural activity across the brain involved in sensory and motor function but also in memory and cognition [48].

**Figure 2:**
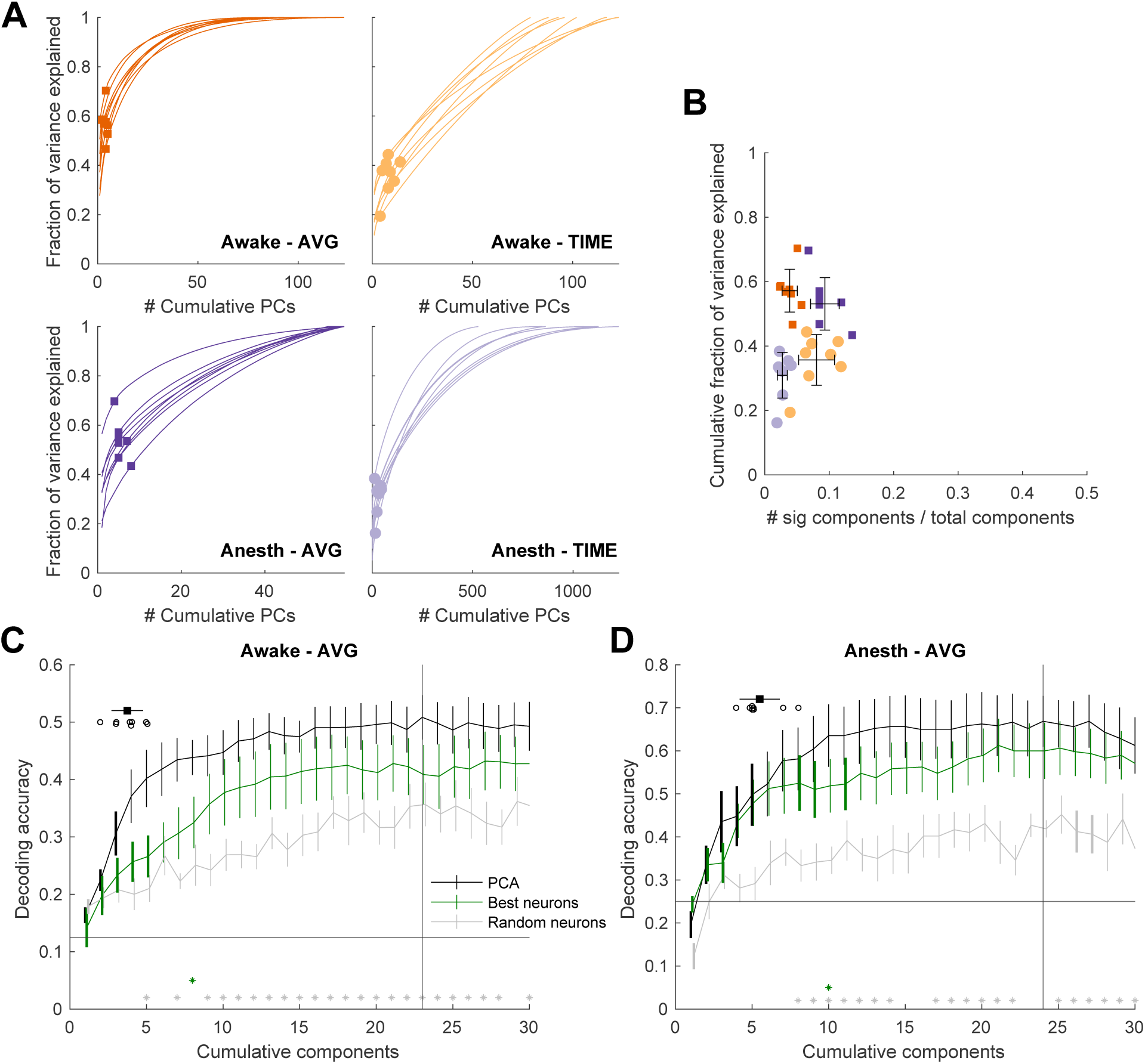
Dimensionality of neural representation of somatosensation in awake and anesthetized animals. **A** Fraction of variance explained by cumulative principal components, calculated using time-varying data concatenated across trials (-TIME) or using responses averaged across time within each trial (-AVG). Similar calculations were made for awake and anesthetized animals (Awake, Anesth). Each line represents the result from a single animal, and each marker (circle or square) denotes the significant number of PCs calculated for that animal. **B** Cumulative proportion of variance explained by the significant PCs for each animal in each dataset, shown as a function of the fraction of significant PCs. Error bars denote mean and standard deviation. Colors as in (A). **C** Limb decoding accuracy in awake mice as a function of cumulative components used (PCs or neurons). Black line shows decoding accuracy from PCs; green line shows decoding accuracy from neurons with the highest average activity across all trials; gray line shows decoding accuracy using randomly-selected neurons. All error bars show mean and standard error. Bold lines denote components in which decoding accuracy was significantly different than the maximum (maximum is identified by the vertical gray line). The black horizontal line signifies decoding performance achievable by chance, and gray vertical line identifies peak decoding performance for PC decoding. Gray stars indicate statistical significance between black (neural PCs) and gray (random neurons) lines, and green stars indicate statistical significance between black and green (best neurons) lines. Black dots in top left show the computed number of significant PCs for each animal (from A), and horizontal error bar in top left shows the mean and standard deviation across animals. A Wilcoxon rank-sum test was performed to compare the score of each component relative to the maximum decoding performance. **D** Analysis as in (C), but for limb identity in anesthetized mice.

Although quantifying the dimensionality of these datasets reveals some consistency of neural encoding structure across behavioral states (anesthetized and awake), this metric does not directly address how much information the neural population encodes about passive limb movements. To address this question, we used a linear classifier (linear discriminant analysis) to determine how well we could decode sensory variables in each dataset as a function of the number of principal components used to describe the data. In both cases, the neural response matrix (from which PCs were derived) was composed of the time-averaged response of each neuron during each trial. In the anesthetized dataset, the decoded sensory variable was the identity of the limb being deflected (contralateral forelimb, ipsilateral forelimb, contralateral hindlimb, ipsilateral hindlimb). In the awake dataset, the decoded sensory variable was the direction of passive movement during center-out trials (one of the eight angles shown in Figure 1B). Both sensory variables could be decoded with modest accuracy. Notably, decoding accuracy plateaued (failed to significantly improve with additional components) at around the average number of significant components found across animals (approximately four; Figure 2C,D). As a control, we compared how well the decoder could perform when trained on the responses of an equivalent number of neurons with the highest average activity across all time, or an equivalent number of randomly-chosen neurons. Although we failed to find a significant improvement in decoding accuracy for PCs vs. the best neurons after correction for multiple comparisons, decoding accuracy continued to improve for the neural decoder beyond the plateau point for PCA. That is, decoding kept improving with increasing numbers of neurons included in the decoder beyond the point that the best decoding performance was reached using PCs. This suggests that information is more efficiently encoded in PC space than in the space of even the most active single neurons. Furthermore, decoding with PCs was significantly better than decoding with randomly-chosen neurons for majority of the cumulative components. These results show that a small fraction of principal components include sufficient information to encode each sensory variable, which aligns with the small fraction of the significant components that explain a large part of the variance in the data. The results support our assertion that neural responses to somatosensation are low-dimensional.

### Joint angle vs. movement vector encoding of limb movements

The variability in single-neuron tuning to passive limb movements across distinct behavioral tasks has yielded mixed conclusions regarding the sensory variables encoded by somatosensory cortex [1, 13, 15, 20–22], including whether somatosensory cortex encodes limb movement in terms of joint angles or as a movement vector. Joint angles refers to the specific configuration of a limb: the angle between the trunk of the body and the upper limb (the shoulder) and the angle between the upper limb and the forelimb (the elbow). In contrast, a movement vector represents the distance and direction between the start and end points of a movement – information that could be abstracted from a specific change in joint angles during a reach. While the two variables are closely related, an identical movement vector can be achieved by distinct changes in joint angles. The awake open-source dataset was elegantly designed to test this hypothesis: the mouse’s forelimb was moved through the same movement vectors with two distinct starting points, generating either center-out or out-center movements (Figure 1B). They found that many single neurons often had similar responses to the same movement vector, regardless of starting point [1]. However, given the variability across single-neuron responses, it is important to additionally characterize population-level responses, including a direct comparison to the joint angle encoding hypothesis. To test the hypothesis that neural populations encode movement vectors vs. joint angles, we used neural responses in the reference frame of PCs of the data to analyze population-level encoding of limb movements in the awake mice.

Each hypothesis (joint-angle or movement-vector) makes a distinct prediction about the structure of neural responses. If the movement vector hypothesis were true, the neural representations of pairs of movement vectors (during inwards and outwards movements, which require different changes in joint angles) would be more similar than neural representations of movements in neighboring directions that had the same starting point (which require similar changes in joint angles). We tested this idea in two frames of reference: neural space and PC space. “Neural space” refers to the set of simultaneously recorded z-scored neural responses to limb movement, where each neuron defines a dimension in neural space. “PC space” refers to the neural response matrix projected onto the set of significant PCs. To make a fair comparison between hypotheses, we projected the neural responses to both inwards and outwards movements into a unified PC space (Figure 3A). We then quantified the similarity between neural representations of movement pairs (joint angle pairs or movement vector pairs) by calculating the correlation coefficient between trial-averaged neural trajectories in PC space (Figure 3B) and neural space (Figure 3C). As a control, we also computed the similarity between each movement and its direct opposite (movements offset by 180*°*), which are dissimilar in terms of both changes in joint angle and in terms of movement vectors. Thus, the 180*°*offset provides a lower bound for similarity in this dataset. We restricted the analysis to datasets from animals with at least 75 simultaneously-recorded neurons. This threshold was chosen from the results of a subsequent analysis, examining neural encoding of limb movements as a function of the number of neurons included in PCA (Figure 3D,E). However, we did not exclude neurons from analysis on the basis of particular responses properties (e.g., tuning to movements).

**Figure 3:**
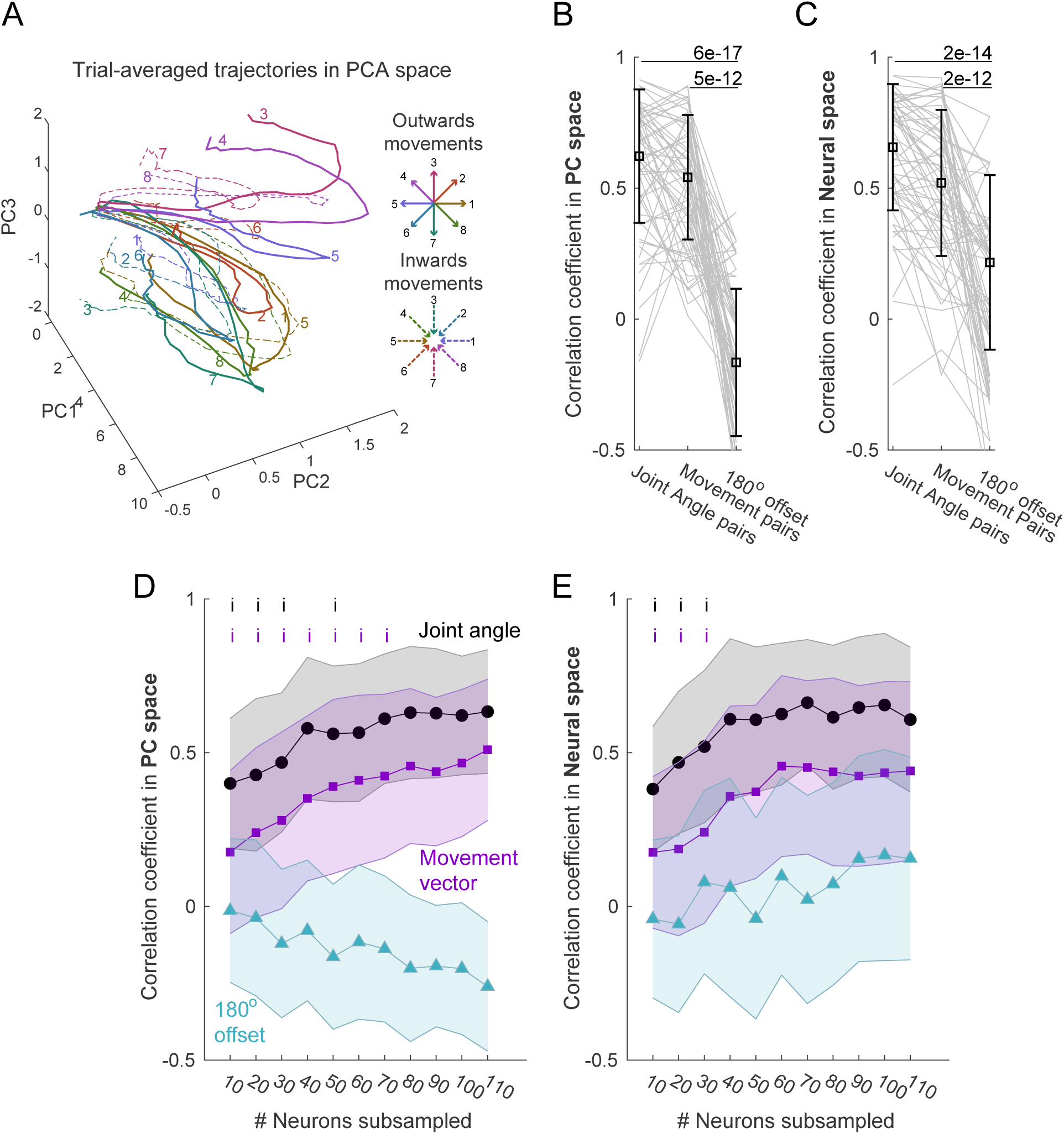
Population-level encoding of passive limb movements in neural and PC spaces. **A** Trial-averaged neural re-sponses to each movement direction during outwards and inwards movements projected onto PC space. Responses are shown along the first three PCs. Colors match movement vector pairs between the outwards and inward movements. **B** Correlation between neural trajectories corresponding to pairs of movements along the first five PCs. Error bars show median and standard deviation. Significant differences between joint angle and movement vector correlations is calculated using a median test. **C** Analysis as in (B), using z-scored neural responses instead of PCs. **D** Analysis as in (B), repeated using sub-samples of the total neural population to compute PCs (restricted to animals with at least 110 neurons recorded simultaneously, N = 3). Black *i* markers indicate stages of improvement in encoding joint angles, defined as a significant difference (p *≤* 0.01, Median test with a Holm-Bonferroni correction for multiple comparisons) from the value calculated at 110 neurons. Purple *i* markers show the same calculation for movement vector encoding. The median correlation coefficients between joint angle encoding, movement vector encoding, and 180° offset pairs are significant at each point (p *≤* 0.01, Median test with a Holm-Bonferroni correction for multiple comparisons). **E** Analysis as in (C), using sub-samples as described in (D) from z-scored neural responses. Colors and statistics as described in (D).

We found that population-level activity in both PC and neural space was encoding both joint angles and movement vectors (Figure 3B,C). In both contexts, we found no statistical difference in the median correlation between joint angle and movement vector pairs. This suggests that neural responses in sensorimotor cortex, when organized within both neural and population-level reference frames, encode multiple variables that describe limb movements. Furthermore, there was no difference in population-level encoding of either joint angles or movement vector pairs between PC and neural spaces (p≥ 0.05, Median Test). The equivalency in encoding results between neural and PC spaces leads us to hypothesize that movement vector information may be extracted from a population of neurons that individually encode joint angles by taking a weighted average of individual neuron activity, which is what happens when projecting neural activity into PC space. This weighted average may marginalize out joint angle information, keeping the variable that was best described by correlated activity across the population: the movement vector.

Using a subset of mice in which 110 neurons were recorded simultaneously (n = 3), we tested this idea by sub-sampling groups of neurons from the entire population (10 repetitions per animal per condition), computing PCs on the smaller neural population, and calculating the correlation between neural trajectories in each subspace. In support of the hypothesis that large populations allow for better abstraction of sensory variables, we found that neural encoding of movement vectors in PC space improved with the inclusion of more neurons in the population beyond the point that encoding of joint angles improved (30-50 neurons for joint angles vs. 70 neurons for movement vectors; Figure 3D). Furthermore, in PC space, encoding of joint angles and movement vectors differentiate relative to encoding of opposing movements (180*°* offset). Interestingly, although larger neural populations can better encode both joint angle and vector-based movements compared to smaller populations, the opposite relationship is observed for encoding of opposing movements in PC space (Figure 3D). In neural space, a similar improvement in encoding for both joint angles and movement vectors is observed as more neurons are included in the population (Figure 3E). However, the encoding of joint angles and movement vector pairs both plateau earlier in neural space, at 30 neurons. In conclusion, our population-level analysis demonstrates that limb movements defined by both joint angles and movement vectors are better encoded by large populations in mouse sensorimotor cortex, supporting prior inferences from analysis of single neurons [1].

### Orthogonality between ipsilateral and contralateral limb representations

The structure of neural representations within motor cortex extends beyond low dimensionality of neural responses. For example, neural representations of both ipsilateral and contralateral limbs in motor cortex co-exist within a single cortical hemisphere yet occupy distinct and orthogonal subspaces [37]. To examine if this same organization exists for somatosensory information from the ipsilateral and contralateral limbs, we used PCA to define a low-dimensional space for neural responses to passive limb movements of the anesthetized mice. We optimized the PCs of neural responses by conducting PCA on passive limb movement trials grouped by contralateral and ipsilateral limbs. The average neural trajectories demonstrate the PCs that best capture variance in neural responses to the contralateral limb movements do a poor job of explaining neural activity related to movement of ipsilateral limbs, and vice versa (Figure 4A,B). Within these specialized subspaces, we quantified the fraction of variance explained by ipsilateral and contralateral responses as a function of cumulative principal components (Figure 4C). We found that, when projected into ipsilateral space, only a small number of PC components are needed to explain a large fraction of the ipsilateral response variance. However, when contralateral responses are projected into ipsilateral space, significantly more PC components are needed to achieve similar explained variance values. We observed the opposite relationship when projecting responses into contralateral space. Therefore, the neural rep-resentation of contralateral and ipsilateral limbs within sensorimotor cortex is orthogonal to each other, analogous to encoding of motor commands.

**Figure 4:**
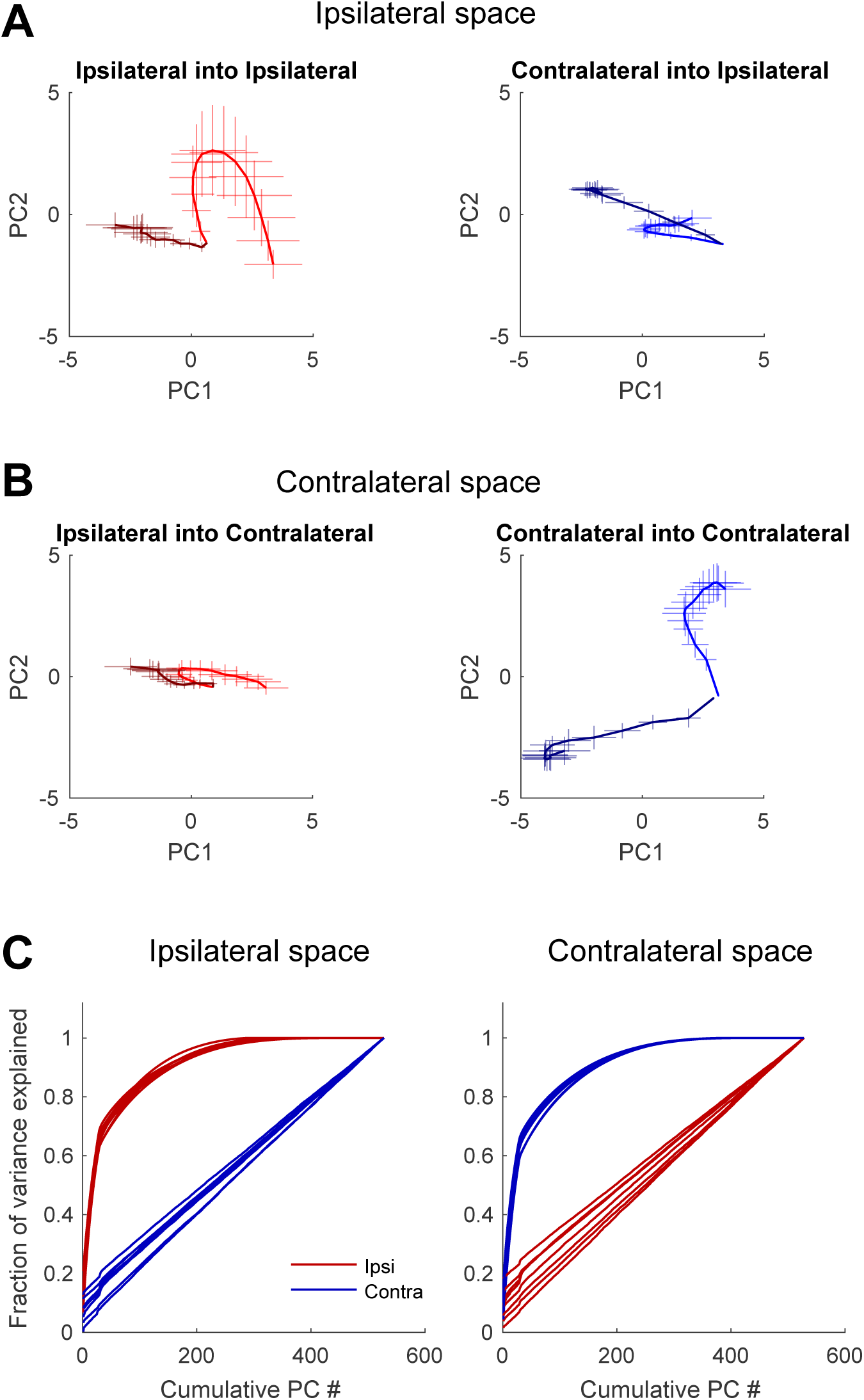
Orthogonality between the projections of ipsilateral and contralateral and forelimb and hindlimb represen-tations in sensorimotor cortex. **A** Across-trial mean and standard deviation of neural trajectories during single-trial limb deflections, projected onto the PC space generated from ipsilateral trial data. Left: ipsilateral trials projected onto PCs obtained from ipsilateral trial responses. Right: contralateral trials projected onto PCs obtained from ipsilateral trial responses. Both examples generated from one anesthetized mouse. **B** As in (A), but projected onto the PC space generated from contralateral trial data. Both examples generated from the same mouse in (A). **C** Cumulative variance of neural responses explained by varying numbers of PCs in *Contralateral* and *Ipsilateral* PC spaces. Each line depicts data from one mouse.

### Conserved representations across animals

A final question is to determine whether the low-dimensional representation of somatosensation is conserved across animals. Within the motor cortex, neural dynamics underlying motor control are well conserved within an animal across time (robust to changes in the recorded neural population), across animals performing the same task, and even across species performing similar behaviors [41, 49, 50]. To determine if this conservation holds true in the sensorimotor system, we used canonical correlation analysis (CCA) to map the trial-averaged neural trajectories from pairs of animals into the same subspace [26, 41, 49]. In anesthetized animals, we analyzed the degree of overlap between neural representations of distinct limbs; in awake animals, we analyzed the degree of overlap between movements of a single limb in different directions. Directly comparing neural encoding of separate limb movements across the anesthetized animals or different directions of a limb across the awake animals is not possible on a trial-to-trial basis due to the variability in limb movements. To compare across animals, we therefore averaged neural trajectories across all trials associated with deflection of a single limb for the anesthetized animals or a direction for the awake animals to obtain separate representative trajectories for different limbs (Figure 5A) or directions within the low dimensional space. CCA was then performed on the matched neural response matrices across pairs of animals. Given two matrices (here, the low-dimensional subspaces associated with somatosensation from two animals), CCA identifies a set of linear transformations that maximally correlate those matrices. Then, the normalized dot product between matched trajectories within this new subspace was used to quantify the degree of matching between animal pairs.

**Figure 5:**
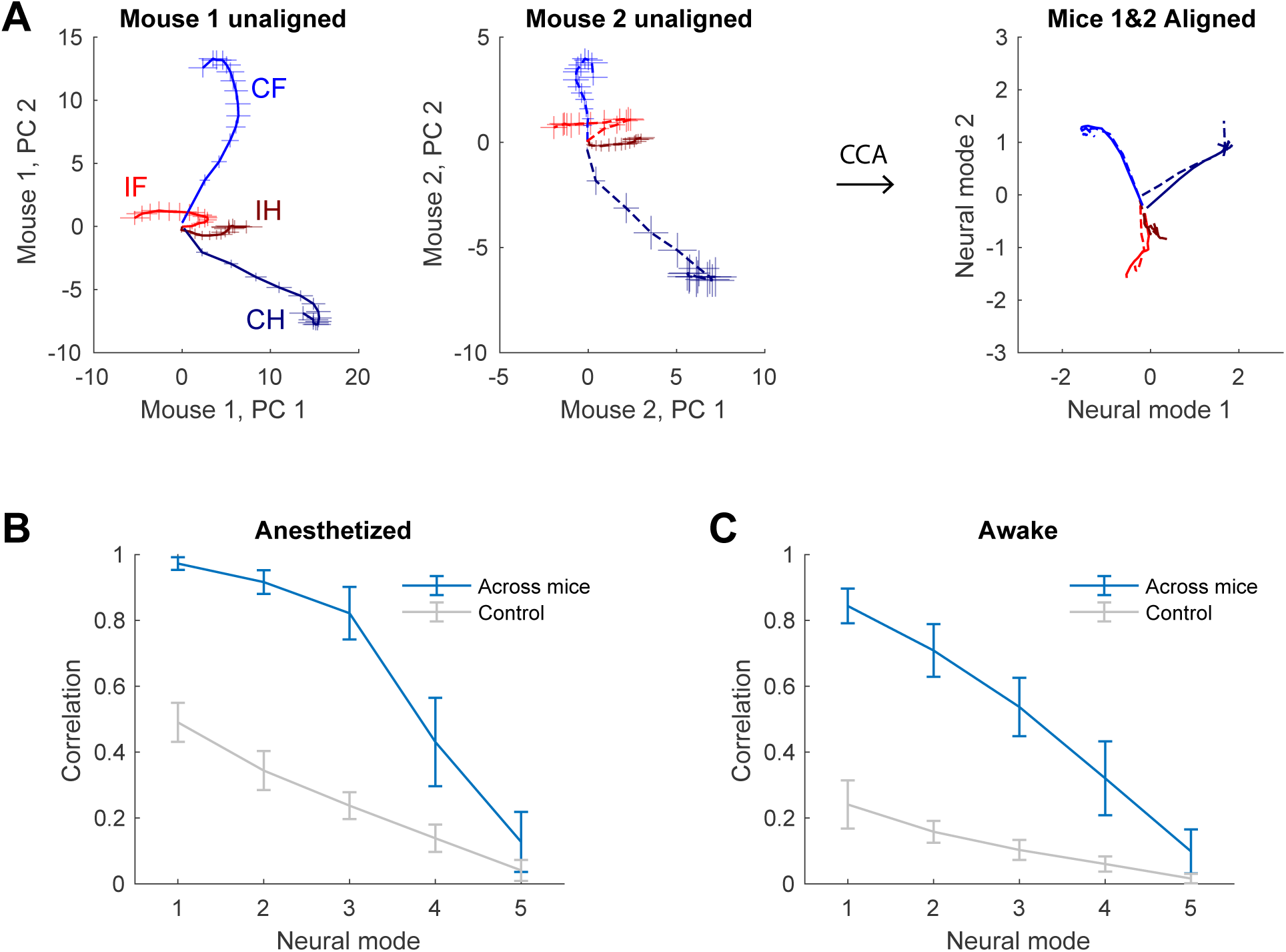
Conservation of the structure of neural representations of limb somatosensation across animals. **A.** Neural trajectories associated with mean single-limb deflections in two sample anesthetized mice and subsequent aligned representations following CCA. Error bars signify standard error across trials. CH: contralateral hindlimb, CF: contralateral forelimb, IH: ipsilateral hindlimb, IF: ipsilateral forelimb. **B.** Correlation coefficient between each neural mode following CCA alignment of average single-limb neural trajectories in 5-dimensional PC space. Bars show mean and standard deviation across subjects (N=8). The control condition shows alignment between true neural data and “average trajectories” computed from a randomly permuted data matrix. Neural modes refer to the set of axes identified by CCA along which data from two animals are best aligned.

We found that the low-dimensional representations of somatosensation are well conserved across both awake and anesthetized animals (Figure 5). As a control, we also performed CCA between true and scrambled datasets; the latter manipulation breaks both the correlations between neurons and the temporal relationship of the dataset but maintains average activity levels. The alignment with scrambled datasets was significantly lower than the true datasets (Figure 5B,C), which shows that CCA cannot align arbitrary pairs of matrices. However, note that increasing the number of dimensions used for alignment can improve alignment, even for arbitrary matrices (Figure S1).

## DISCUSSION

The purpose of this manuscript was to investigate the structure of neural activity within mouse sensorimotor cortex as it relates to somatosensory inputs from the limbs. Our major idea was to “summarize” activity within a large population of neurons by using principal component analysis – a standard form of dimensionality reduction that identifies correlated activity across independent variables (here, across neurons). Using these low-dimensional representations, we were able to establish that the structure of neural activity in mouse sensorimotor cortex mirrors the observed low-dimensional encoding of motor commands and is well preserved across animals (Figures 2,5), including maintaining orthogonal representations of ipsilateral and contralateral limbs (Figure 4). This population-level approach additionally yielded insight into the way that neural populations encode variables that differ from single neuron encoding. We found that an abstract representation of limb movements (a movement vector) can emerge from a population of neurons (Figure 3). Together, our results illustrate how the structure of neural representations in sensorimotor cortex may facilitate sensorimotor integration across the brain.

### Somatosensation includes both touch and proprioception

Two major classes of sensory inputs that are involved in somatosensation, touch and proprioception, both activate neural responses in overlapping brain regions [1]. Within the somatosensory cortex, touch and proprioception recruit distinct populations of neurons such that blocking tactile inputs does not significantly alter the responses of proprioceptive-encoding neurons to passive limb movements [1]. Therefore, tactile sensations induced during the course of the experiments described here (primarily pressure upon the paw(s) from the experimenter or induced by the animal’s grasp of the robotic manipulandum) would be expected to activate separate subpopulations of recorded neurons that could potentially increase the total dimensionality of neural activity. While touch has the capacity to be a high-dimensional signal in the sense that higher PCs can be used to accurately identify the texture presented to an animal [51], we found in this current study that neural activity in responses to somatosensory inputs were low-dimensional. Notably, even when touch is considered high-dimensional (as in primate neural encoding of texture), the first two PCs explain approximately 80% of the variability in neural responses to textures [51]. In this context, finer details about the texture are present in higher PCs. In our current experiment, tactile inputs are relatively constant across trials and across limbs. This consistency may explain our results that neural activity patterns across neurons are correlated and can be well summarized by a relatively small number of PCs relative to the size of the simultaneously recorded neural population. However, there is some evidence that there are distinct neural mechanisms for encoding texture in the mouse and in the primate. In the study referenced above [51], the first PC was most strongly correlated with the roughness of a texture, which dominated the time-varying changes in neural activity patterns. In contrast, neural activity in mouse somatosensory cortex isn’t time-locked to vibration frequency (which could encode roughness), but instead neurons are tuned to specific input frequencies [52]. This encoding mechanism could increase the dimensionality of the touch signal in mouse cortex, but the datasets analyzed in this current manuscript are not designed to evaluate this possibility.

### The intrinsic dimensionality of somatosensation

*Must* the neural encoding of somatosensation in mouse sensorimotor areas be low-dimensional? We discussed some implications for the dimensionality of touch in the previous section, so we focus here on proprioception. The sense of proprioception is based on sensors embedded in the tendons, muscles, and joints that are active both during static limb configurations and during dynamic changes in posture [5]. Receptor activity conveys redundant information about twenty-two muscles in the mouse forelimb and forty muscles in the hindlimb [53, 54], which form a smaller number of functional groups that enact coordinated activity across multiple muscles. For example, in the mouse hindlimb, muscles coordinate to rotate, flex, or extend the knee, hip, and ankle [54]. At an even more abstract level, limb posture and movement can be represented simply as a set of joint angles (of the hip, knee, and ankle), changes in which are highly correlated across limb movements [55]. Movement can also be represented as a vector that, unlike joint angles, is independent of the starting and end points of the reach (e.g., distance and direction). Therefore, there are relatively few independent variables to encode by the hundreds of thousands of neurons in sensorimotor cortical areas [42]. In consequence, a reasonable hypothesis regarding the dimensionality of somatosensory information within the cortex is that it would be low-dimensional. While we find this to be the case, neural anatomy suggests that the dimensionality of sensory information *could* increase along the cortical hierarchy, as the number of neurons in the cortex is roughly an order of magnitude larger than in the thalamus [42, 43]. This significant anatomical expansion could have mediated an expansion of the dimensionality of somatosensation within sensorimotor cortex [44]. Therefore, the dimensionality of somatosensory information in the cortex is not obvious a priori.

### Robustness of results across datasets

We used two complementary datasets (anesthetized passive multi-limb and awake passive forelimb) to show that somatosensory information from the limbs occupies a low-dimensional subspace within the neural population response, regardless of the behavioral state of the animal. The anesthetized data is valuable because it contains the responses of neurons within mouse sensorimotor cortex to passive deflection of each of the four limbs. These deflections were performed manually, which resulted in high trial-to-trial variability in limb trajectories within an acute range of angles (Figure S2). High variability in limb trajectories is critical to estimating the true dimensionality of somatosensation, as dimensionality is reduced during highly stereotyped movements [56].

Analysis of the awake dataset was essential to validate the hypotheses generated from the anesthetized data, including verifying that the measured dimensionality does not arise simply from changes in network structure during anesthesia [57, 58]. In the awake passive forelimb experiments conducted by another lab [1, 59], mice held a robotic manipulandum during passive movements in both center-out and out-center directions (Figure 1B). Unlike passive movements under anesthesia, here, the mouse’s forelimb was moved over a wide range of angles (in eight directions from 0 to 360 degrees). However, given the robotic control of the movements, limb movements in any particular direction were highly stereotyped, which might artificially depress the dimensionality of neural responses [56]. The variability in experimental conditions across anesthetized and awake datasets ensures the robustness of our conclusions, as we see a similar relationship between the fraction of significant PCs to the cumulative proportion of variance explained across datasets (Figure 2).

### Movement vectors vs. joint angles

We found that neural activity patterns represented movement vectors as well as joint angles, and that encoding of both variables improves when the principal components were calculated based on the activity of a large population of neurons (Figure 3). Why should somatosensory inputs be represented in the form of a movement vector rather than more concretely as changes in joint angles? One possibility, suggested by computational modeling, is that in many sensorimotor tasks, *“what”* needs to be done is invariant to *“where”* it is being done in space [60]. This “what” representation has been observed in higher brain areas, including the parietal and premotor cortices, which tend to encode movement intention rather than limb kinematics [61–63]. Furthermore, premotor cortex is known to convey movement intention (efference copy) to somatosensory cortex [64]. Therefore, sensorimotor cortex may abstract changes of joint angles into vectorial representations of movement to efficiently communicate with higher brain areas.

Human and animal perception seems to better align with higher-level descriptions of movements [1, 65, 66]. For example, mice are more likely classify passive limb movements according to the direction of movement relative to the body (towards vs. away) rather than by the changes in joint angles elicited by that movement [1], and humans are more precise at aligning a cursor with their hand position than at rotating a line to match a joint angle [66]. Furthermore, electrical microstimulation of the human parietal cortex elicits proprioceptive sensations that describe the direction of movement (upwards, forward, etc. [65]), which emphasizes that our conscious experience of movement is its direction.

### Implications for clinical neural prostheses

Our population-level analysis of neural activity in sensorimotor cortex of mice has shown that the encoding of somatosensation mirrors that found in other cortical areas involved in sensorimotor processing, and that this similarity could facilitate the integration of information across brain areas. This basic scientific result has important implications for neural prostheses, as somatosensory cortex is seen as a key anatomical target for neuromodulation to provide artificial sensation in bidirectional brain-machine interfaces [67–72]. Understanding how somatosensation is encoded in sensorimotor cortex will help us design optimal artificial sensory signals that will integrate with natural sensation and ultimately improve the performance and embodiment of neural prostheses [73–79].

## METHODS

### Mice

All procedures were approved by the Institutional Animal Care and Use Committee at Purdue University. Eight CaMKII-tTA (JAX stock #007004) x TRE-GCaMP6s (JAX stock #024742) mice aged 3-6 months were used for 2-photon calcium imaging. Mice were housed in groups of no more than five per cage in a temperature and humidity-controlled room on a 12-hour light/dark cycle with *ad libitum* access to food and water.

### Surgical Procedures

For all surgical procedures, mice were anesthetized with isoflurane at 3% for induction and maintained between 1.5-2.5% to areflexia. All mice were first surgically implanted with a custom titanium headplate (8 mm circular inner diameter) placed over the right cortical hemisphere and secured with dental cement. Allowing at least three days for recovery, a 5 mm craniectomy was then made over the primary somatosensory and primary motor cortical areas (stereotaxic anatomical coordinates determined from Paxinos & Franklin). A 5 mm round glass window was placed over the brain and secured with a cyanoacrylate mixture. Mice were then given at least three days to recover from surgery before undergoing imaging.

### Anesthetized Experiment

Mice were first anesthetized with isoflurane at 3% for induction and maintained at 0.9-1.1% to preserve neural activity during recordings. A heating pad maintained at 37°C (World Precision Instruments) was placed under the mice for the duration of anesthesia. Mice were carefully examined throughout the duration of the experiment for respiratory pattern and areflexia to ensure a consistent level of anesthesia. The mouse’s headplate was secured to a custom headplate holder to fix the head for two-photon imaging (Neurolabware microscope, Scanbox software). Limb movement during the experiment was video recorded at 30 fps (The Imaging Source).

Two-photon imaging in 1x magnification (approx. 1.9 x 1.2 mm FOV), image sampled at 7.8 Hz, was used to record neural activity patterns in sensorimotor cortex during passive limb movements. The imaging windows were positioned in various locations over sensorimotor cortex (Figure S3) to obtain neural recordings across the entire region. Passive limb movement was performed by manually brushing the entire limb, starting at an area of the limb close to the trunk and moving towards the paw. This was done to each limb five times for three cycles, with a total of 15 passive movements for each limb. The order of limb movement was randomized for each mouse, and ample time (∼10 seconds) was given between each movement to allow the fluorescent signal to return to baseline levels before the next movement.

Two-photon images were processed using Suite2p [80], an image processing pipeline that performs image registration, cell detection, and signal extraction from detected cells. Extracted fluorescence traces were processed and neuropil signals were removed by subtracting neuropil signals multiplied by 0.7 and standardized (z-scored).

### Principal component analysis (PCA) and dimensionality analysis

PCA was used in this manuscript as a method of dimensionality reduction. The goal of PCA is to identify a new set of axes, composed of a linear combination of the original dataset, that captures correlations between independent variables such as neurons. PCA was performed using the MATLAB *pca* function in the stats toolbox.

For the anesthetized and awake passive movement datasets, multiple trials were conducted within each recording session for each animal. Prior to PCA, single-trial responses were extracted from the total recorded session. The response of a single neuron to passive movement on a single trial was taken relative to the baseline fluorescence (the value of the neural activity just prior to movement onset). For anesthetized mice, trial onset and offset was defined as the neural imaging frame on which movement began and ended, which was manually identified from the concurrent 30 fps video recordings during the experiments. For awake mice, trial onset and offset times were as defined as in [1] and computed using the provided open-source code [59].

The total neural response of a population of neurons to a set of trials was found to be a tensor of size [the number of trials, the number of time points per trial, the number of neurons]. As described above, in the case of the anesthetized data, the number of timepoints per trial could vary. We used PCA to compute the dimensionality of the neural responses under two conditions. First, we averaged single-neurons responses within each trial ([trials, neurons]; *ANES-AVG* and *AWAKE-AVG* in Figure 2), which yielded a single value per trial. Second, we used the time-varying responses during the trial, concatenating the data across trials ([trials×time points, neurons]; *ANES-TIME* and *AWAKE-TIME* in Figure 2). The first strategy results in projections of shape [trials, total PCA components] and the second results in projections of shape [trials×time points, total PCA components]. The total number of PCA components that can be found is constrained by the smallest dimension in these matrices, i.e. *min(trials, neurons)* or *min(trials*×*time points, neurons)*. To determine the number of significant PC computed from the neural data, we used the Marchenko-Pastur law, which compares the singular values of the data correlation matrix to a theoretical maximum achievable by chance [46]. The shape of the datasets, the shape of the projections, the number of significant components, and the variance explained by the significant components for each mouse are specified in Tables S1-2.

### Decoding limb movement from neural activity patterns

For both anesthetized and awake mice, we used the average responses of single neurons on single-trials to decode which limb was moved (anesthetized) or which direction the limb was passively deflected (awake). The average response was calculated as the mean activity over the 2 seconds following trial onset. These average responses were used to construct a neural response matrix across all experimental trials, X, of size [# trials, # neurons]. The principal components, W, of that matrix were found, and the neural responses were projected into principal component space, Z = XW. We used linear discriminant analysis (MATLAB functions *fitcdiscr* and *predict* as a decoder, and leave-one-out cross validation (LOOCV) to evaluate the prediction accuracy. Under LOOCV, the model is trained on all of the data point but one, and is tested on the remaining data point.

The experimental question was how well we could decode with increasing numbers of principal components. Therefore, the decoding analysis was first performed using just the first column of Z (PC1), then the first two columns (PCs 1 and 2), and so on. Decoding accuracy is plotted as a function of the cumulative number of PCs used. The same analysis was repeated using single neuron responses. In one version, we used the neurons with the largest average response amplitudes across all trials, starting with the most responsive neuron. In a second control, we chose random neurons for the decoder (random columns of X). In this version, a new random selection was made for each of the cumulative component values. That is, neuron “1” might be used for decoding from a group of two neurons, but might not be selected when decoding from a group of ten neurons.

### Neural encoding of joint angles vs. movement vectors

This analysis was conducted on neural data from [1], using their provided MATLAB code to extract neural responses to single trials. *Neural space:* A three-dimensional neural response tensor was found for each mouse [trials, timepoints per trial, neurons]. Trials included successfully completed center-out and out-center movements (Figure 1B). Finally, a trial-averaged neural response trajectory was found for each movement in each of the outwards and inward directions. *PC space:* The neural response tensor was converted into a two-dimensional matrix of size [trials× timepoints per trial, neurons]. The PCs of this matrix were then found using the MATLAB function *pca*, and the original neural response tensor was projected onto PCs 2-8 (where 8 is approximately the average number of significant components found for individual mice in this dataset). PC1 was excluded as it tends to be the average response of all neurons. This formed the PC space. Finally, a trial-averaged neural response trajectory was found for each movement in each of the outwards and inward directions.

To determine whether the neural population activity best encoded joint angles or movement vectors, we computed the correlation between pairs of neural trajectories in both *neural space* and *PC space*. To accurately parallel the PC space that excluded PC1, we also subtracted the population-average response across all trials from the single-neuron response per trial in calculating the “neural space” responses. Correlation was calculated as a dot product between trial-averaged neural response vectors 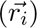, divided by the length (norm) of each vector: 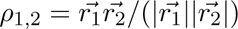. To measure similarity between movement vector pairs, we calculated the correlations between matched center-out and out-center movements. To measure similarity between joint angle pairs, we used the minimum of the correlations between the neural response to center-out movement i and the neural response to center-out movement i − 1 and i + 1.

To examine the relationship between neural encoding and neural population size, we identified a subset of mice (n = 3) in which at least 110 neurons were imaged simultaneously. We repeated the process described above using subsample of the entire neural population (sampling with replacement from each mouse, 10 repetitions at each size).

### Orthogonality between the projections of ipsilateral and contralateral limb representations

Two different PC spaces were obtained using the neural responses of the anesthetized animals for contralateral limbs and ipsilateral limbs, separately (Figure 4). The variances explained of the contralateral responses projected onto the contralateral and ipsilateral spaces were both normalized by the sum of the variance explained of the contralateral responses projected onto the contralateral space because it is the available maximum. Similarly, the variances explained of the ipsilateral responses projected onto the contralateral and ipsilateral spaces were both normalized by the sum of the variance explained of the ipsilateral responses projected onto the ipsilateral space.

### Conserved representations across animals

To test for the similarity of neural representations of limbs movements across animals, we used CCA to align the average neural trajectories associated with movement of each limb (anesthetized mice) or associated with each movement direction (center-out passive movements in awake mice). For the anesthetized dataset, trials were of variable length, and the entire trial was used to compute the PCs of the neural responses. However, to compute the average neural trajectory associated with each limb movement, neural activity in each trial was resampled to acquire the response over the first two seconds (15 samples at 7.8 Hz sampling rate) following movement onset [trials, 15 timepoints, neurons]. A duration of 15 samples was chosen as it was the mode of the unaltered trial durations. This neural response matrix was permuted to yield a matrix of size [trials×15, neurons] and was then projected onto the first three PCs (within the lower end of range of significant components of this type of dataset for the animals). After permuting the matrix once more into a tensor [trials, 15 timepoints, 3 PCs], the trials associated with movement of each limb were averaged to generate a matrix of average responses to passive deflection of each limb [15 timepoints, 3 PCs]. This process was repeated for each mouse, yielding eight neural response matrices. Then, we used the MATLAB function *canoncorr* to map neural response matrices from pairs of mice into a common reference frame. Once the data was in the same reference frame, we calculated the correlation between the neural trajectories associated with each limb from each mouse (using the same equation described above). As a control, the original data matrix was shuffled [trials×timepoints, neurons], over both the rows and columns, and the true data from one mouse in each pair was mapped to the shuffled representation. This calculation sets a baseline level for how well CCA can match two matrices when one of them is meaningless (random). Note, however, that the correlations depend on the number of PCs that are chosen. As this number grows, arbitrary matrices can be flexibly mapped to one another. The same analysis was repeated for the awake dataset, but as the trials were of equal lengths, there was no need to resample neural data.

## ACKNOWLEDGMENTS

This work was supported by the National Science Foundation as part of the HDR Institute: Accelerated AI Algorithms for Data-Driven Discovery (award #2117997). We thank Elizabeth Frazier for providing valuable feedback on the analyses in this manuscript.

## AUTHOR CONTRIBUTIONS

M.H.L. and M.C.D. designed the experiments, M.H.L. conducted the experiments, M.H.L., S.P., and M.C.D. designed the analysis, and S.P. and M.C.D. conducted the analysis.

## AUTHOR DECLARATION

The authors have no competing interests to declare.

---

**Table S 1:** The shape of the data (Data shape), the shape of the projected data on principal component space (PC projection shape), the obtained significant component (Significant component), and the cumulative proportion of variance explained by the significant component (Variance explained) of the anesthetized mice. num trial: the number of trials; num neuron: the number of neurons; len seq: the length of the concatenated sequence; num sample: the number of sample used for principal component analysis; num pc: the total number of the principal components.

| Data type | Animal name | Data shape | PC projection shape | Significant component | Variance explained |
| --- | --- | --- | --- | --- | --- |
| Anesthetized mouse, Average | - | (num_trial, num_neuron) | (num_sample, num_pc) | - |  |
|  | Mouse1 | (57, 1719) | (57, 57) | 5 | 0.53 |
|  | Mouse2 | (53, 901) | (53, 53) | 5 | 0.47 |
|  | Mouse3 | (60, 1773) | (60, 60) | 7 | 0.54 |
|  | Mouse4 | (60, 2229) | (60, 60) | 8 | 0.43 |
|  | Mouse5 | (60, 1950) | (60, 60) | 5 | 0.47 |
|  | Mouse6 | (60, 1191) | (60, 60) | 5 | 0.57 |
|  | Mouse7 | (60, 528) | (60, 60) | 4 | 0.70 |
|  | Mouse8 | (60, 1302) | (60, 60) | 5 | 0.55 |
| Anesthetized mouse, Concat | - | (len_seq, num_neuron) | (num_sample, num_pc) | - |  |
|  | Mouse1 | (617, 1719) | (617, 617) | 55 | 0.58 |
|  | Mouse2 | (555, 901) | (555, 555) | 49 | 0.59 |
|  | Mouse3 | (829, 1773) | (829, 829) | 57 | 0.61 |
|  | Mouse4 | (421, 2229) | (421, 421) | 57 | 0.62 |
|  | Mouse5 | (723, 1950) | (723, 723) | 60 | 0.60 |
|  | Mouse6 | (688, 1191) | (688, 688) | 58 | 0.60 |
|  | Mouse7 | (720, 528) | (720, 528) | 50 | 0.63 |
|  | Mouse8 | (677, 1302) | (677, 677) | 55 | 0.63 |

**Table S 2:** The shape of the data (Data shape), the shape of the projected data on principal component space (PC projection shape), the obtained significant component (Significant component), and the cumulative proportion of variance explained by the significant component (Variance explained) of the awake mice. num trial: the number of trials; num neuron: the number of neurons; len seq: the length of the concatenated sequence; num sample: the number of sample used for principal component analysis; num pc: the total number of the principal components.

| Data type | Animal name | Data shape | PC projection shape | Significant component | Variance explained |
| --- | --- | --- | --- | --- | --- |
| Awake mouse, Average | - | (num_trial, num_neuron) | (num_sample, num_pc) | - |  |
|  | b0039_210525dff | (94, 102) | (94, 94) | 3 | 0.57 |
|  | c0008_201120dff | (126, 123) | (126, 123) | 3 | 0.59 |
|  | c0010_201113dff | (106, 117) | (106, 106) | 4 | 0.58 |
|  | c0021_210310dff | (245, 79) | (245, 79) | 4 | 0.57 |
|  | d0008_201124dff | (165, 123) | (165, 123) | 5 | 0.70 |
|  | d0010_201113dff | (100, 88) | (100, 88) | 5 | 0.56 |
|  | f0010_201120dff | (86, 96) | (86, 86) | 2 | 0.53 |
|  | m0010_201211dff | (121, 93) | (121, 93) | 4 | 0.58 |
| Awake mouse, Concat | - | (len_seq, num_neuron) | (num_sample, num_pc) | - |  |
|  | b0039_210525dff | (3196, 102) | (3196, 102) | 4 | 0.19 |
|  | c0008_201120dff | (4284, 123) | (4284, 123) | 8 | 0.44 |
|  | c0010_201113dff | (3604, 117) | (3604, 117) | 8 | 0.31 |
|  | c0021_210310dff | (8330, 79) | (8330, 79) | 5 | 0.37 |
|  | d0008_201124dff | (5610, 123) | (5610, 123) | 14 | 0.41 |
|  | d0010_201113dff | (3400, 88) | (3400, 88) | 9 | 0.37 |
|  | f0010_201120dff | (2924, 96) | (2924, 96) | 7 | 0.41 |
|  | m0010_201211dff | (4114, 93) | (4114, 93) | 11 | 0.33 |

**Figure S 1:**
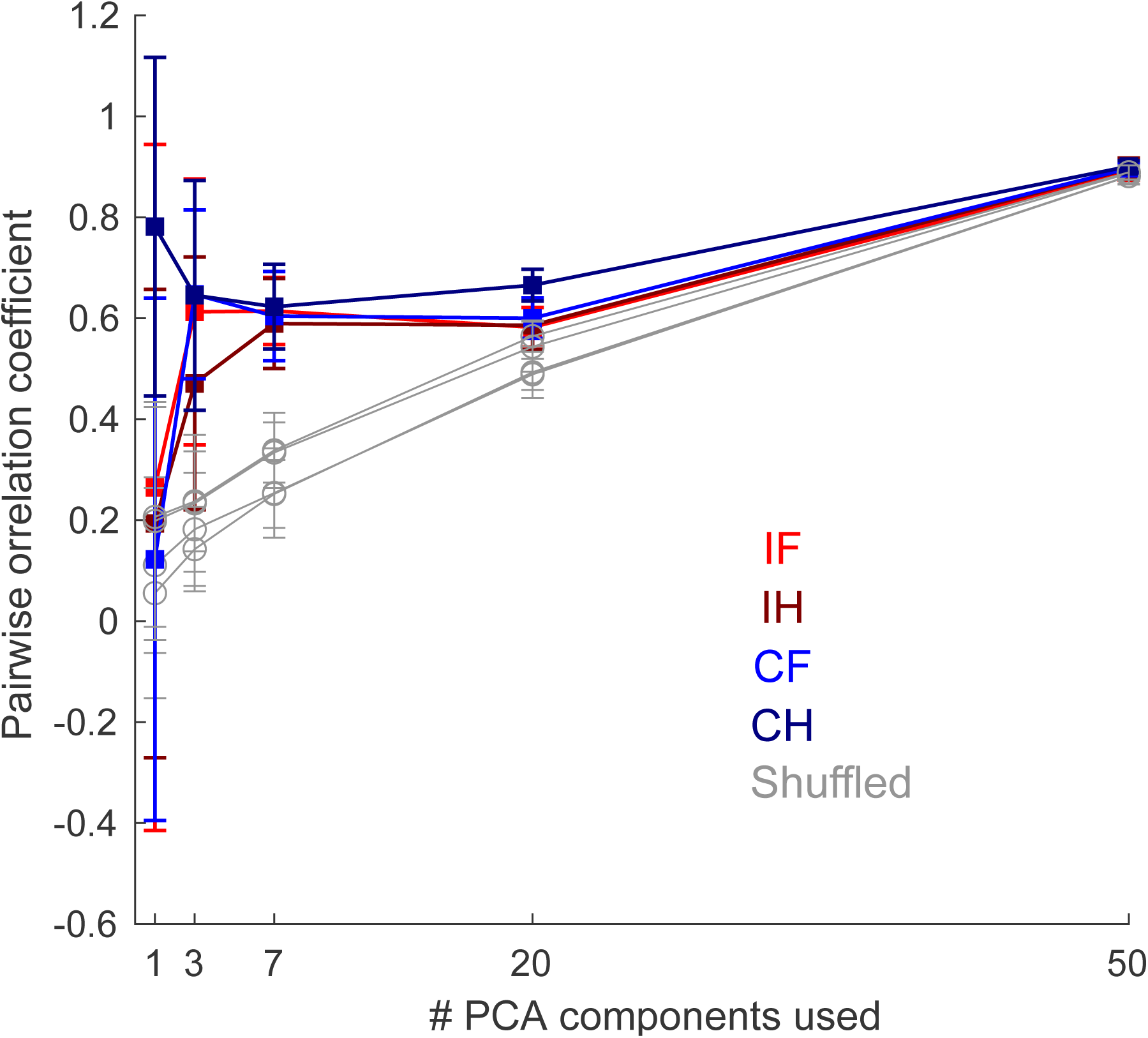
Correlations between aligned neural trajectories as a function of the number of principal components used for alignment.

**Figure S 2:**
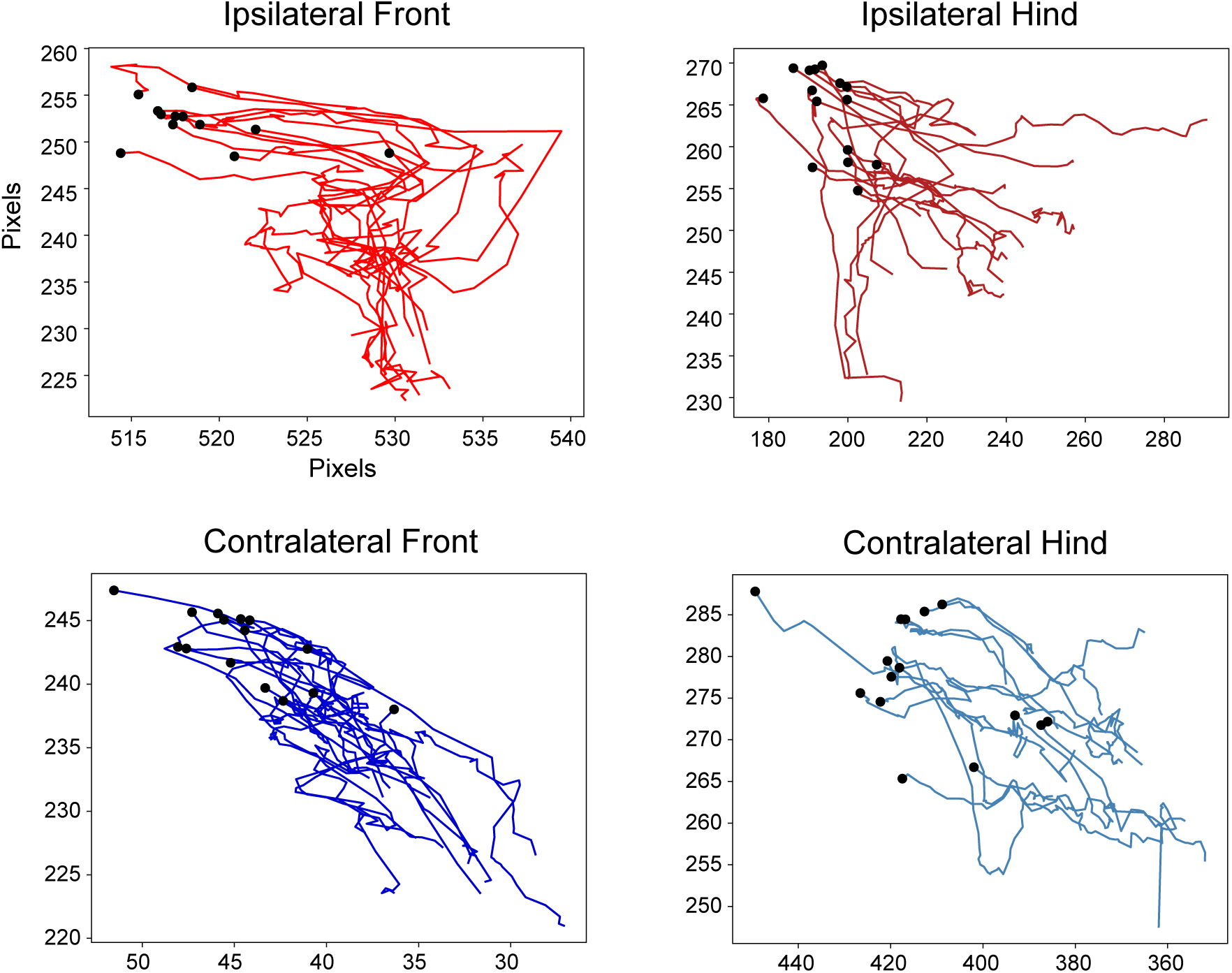
Trajectories of limb end points of Mouse 1 in video frames with the size of 540 × 400 in anesthetized mouse experiment. Black points show the initial points of trials.

**Figure S 3:**
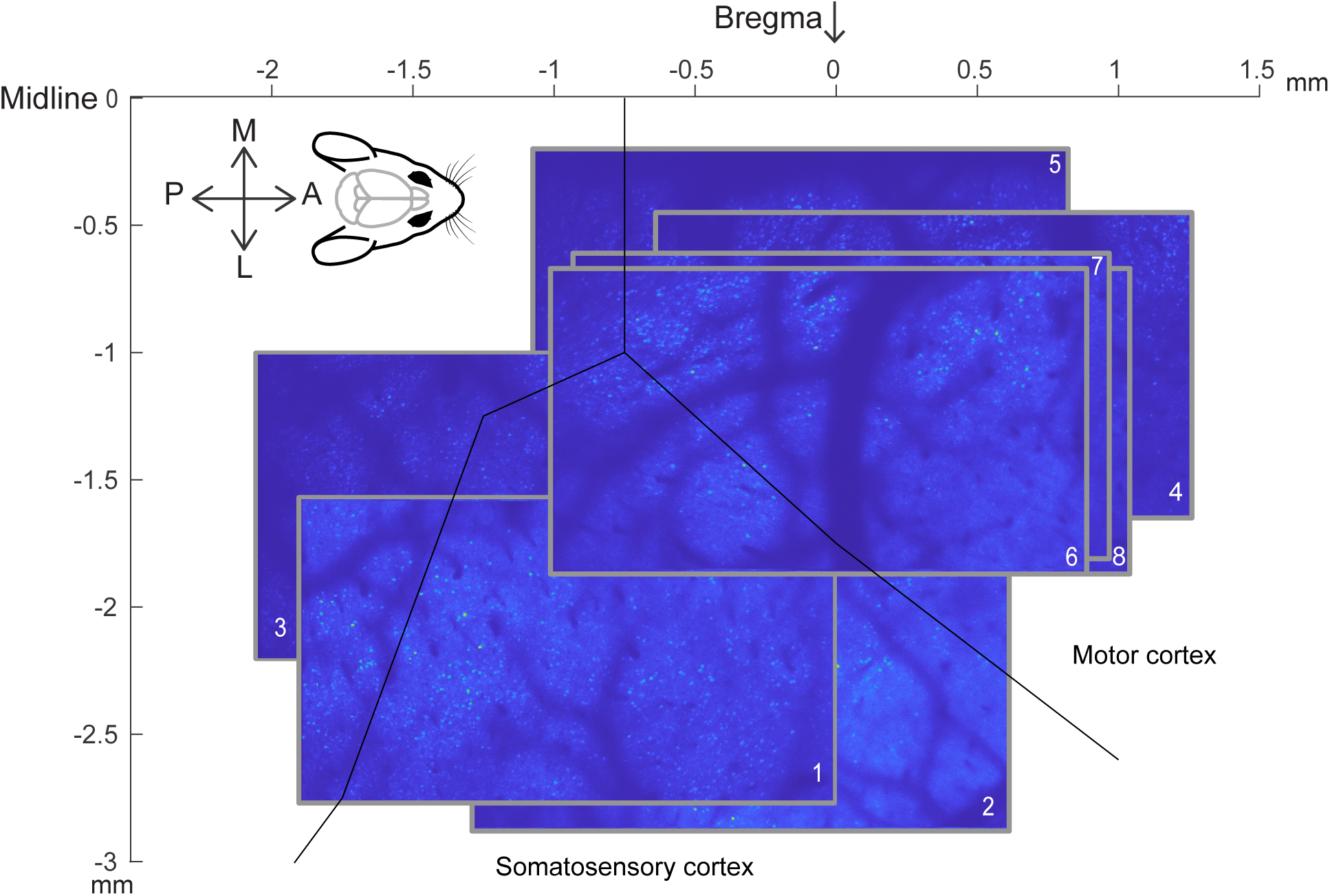
Imaging windows for anesthetized mice (numbered as in the main text and figures, e.g. Mouse 1). The black line denotes putative anatomical demarcations of somatosensory and motor cortices (Krumin et al., 2018).

